# Intragenic Transcription from Defective HIV Proviruses Triggers Interferon Responses in Myeloid Cells

**DOI:** 10.64898/2026.04.06.716685

**Authors:** Jonathan Kilroy, Aparna Deokar, Samantha Patalano, Juan Fuxman Bass, Manish Sagar, Suryaram Gummuluru, Andrew J. Henderson

## Abstract

The persistent HIV-1 reservoir includes a subset of cells harboring transcriptionally repressed latent HIV-1 that contributes to rebound upon antiretroviral treatment (ART) interruption. However, the majority of the reservoir consists of defective proviral genomes with mutations that prevent production of HIV −1 virions. People with HIV (PWH), even with suppression of viremia, demonstrate comorbidities of the central nervous system, heart, gut, and general aging-associated inflammation. Previously, we identified a transcriptionally active element within the envelope gene (*env*) of HIV-1 which mediates the expression of aberrant HIV-1 RNAs. We hypothesize that spurious expression of defective proviruses contributes to the general inflammation that drives these comorbidities. We observed correlations between levels of pro-inflammatory cytokines in serum of PWH and levels of HIV-1 transcripts from this intragenic promoter. To investigate the impact of defective proviruses, we employed CRISPR-Cas9 to render the 5’ Long Terminal Repeat (LTR), which acts as the enhancer and promoter for proviral transcription, non-functional. HIV-1 infected cells harboring this deletion produce significantly higher levels of IP-10 *in vitro* in both monocytic cell lines and primary monocyte-derived macrophages. Transcripts generated from the *env* promoter, include a 5’ cap and polyA tail and the induction of IP-10 expression was dependent on the cytosolic innate immune sensing pathway components MDA5 and MAVS. We propose that defective HIV proviruses contribute to chronic inflammation in PWH via an MDA5-dependent induction of type I interferon pathways.

**Importance:** People with HIV-1 are at higher risk of developing age-associated comorbidities and immune exhaustion even when receiving antiviral treatments and having no detectable viremia. Transcription and translation have been documented from latent and defective proviruses, but their impact on inflammation associated with chronic HIV-1 infection remains poorly understood. The significance of this work is in identifying a role for defective HIV-1 proviruses and correlating their transcription in triggering a type I interferon response. These results highlight the importance of the persistent defective HIV-1 proviruses and understanding their impact on driving chronic inflammation to inform future strategies to assure people with HIV-1 healthy living and aging.

## Introduction

Human Immunodeficiency Virus (HIV) remains a global burden even with the availability of effective antiretroviral therapies (ART) [1]. Recent estimates put the number of people with HIV (PWH) worldwide at approximately 40 million, with an annual new infection rate of 1.3 million per year [2]. Upon infection, HIV-1 integrates a DNA copy of its genome into the host cell’s DNA establishing a provirus that persists for the life of the cell [3, 4]. Transcription of a subset of these proviruses is repressed, and in the context of ART, will evade targeting by the immune response to generate a reservoir of infected cells [5–8]. These latently infected cells, which include CD4+ T cells, macrophages, dendritic cells, and microglia, present the primary barrier to a cure since they support new virus replication and infection following treatment interruption [9–11]. Low levels of cell-associated HIV-1 RNA have been identified in peripheral blood mononuclear cells (PBMCs) from PWH on ART, even when viral load is below the limit of detection [12–14]. While ART prevents the spread of HIV, it is not a cure [15, 16].

PWH are at higher risk of developing comorbidities including neurological deficits, frailty, cardiovascular disease, gut dysbiosis, and a general accelerated aging, often referred to as “inflammaging” [17–20]. This increased susceptibility to inflammatory diseases persists in PWH actively being treated with ART as well as controllers who repress HIV-1 in the absence of ART indicating that these chronic comorbidities are not solely driven by active HIV-1 replication [21–25]. It has been proposed that spurious reactivation of the latent reservoir or blips, ART interruption leading to rebound, and long-term immune cell exhaustion and dysregulation of inflammation contribute to HIV-1 associated inflammation [26, 27].

Recent work has identified multiple pathways that initiate HIV-1 mediated inflammation. For example, CARD8 inflammasome activation by HIV-1 protease and NLRP1 inflammasomes triggered by unspliced HIV-1 RNA, induce the production of IL-1β and pyroptosis in infected cells [28, 29]. In addition, expression of peptides during latent infection have been shown to lead to antigen presentation and shaping of T cell responses [14, 27]. Furthermore, unspliced intron containing HIV-1 RNAs (icRNAs) induce type I interferon responses in myeloid cells through a MAVS dependent pathway [30, 31]. Importantly, the majority of persistent HIV-1 genomes are defective proviruses that harbor deletions, insertions, inversions, mutations, and other errors that prevent the production of new virions, however, these defective proviruses are transcriptionally and translationally active producing HIV-1 RNAs and proteins that potentially perpetuate inflammation [5, 6, 13, 32]. Therefore, HIV-1 generates a variety of pathogen associated molecular patterns (PAMPS) that are detected by pattern recognition receptors (PRRs) to initiate general inflammation [29, 33–37].

We previously identified a transcriptional element within the HIV-1 *env* gene that supports transcription even in the absence of a functional 5’LTR [38]. Transcripts driven by this intragenic HIV-1 promoter have been detected in PBMCs from PWH on ART [26, 38]. We hypothesize that active transcription from defective proviruses induces innate immune responses, and thus contributes to the persistent inflammation and immune dysregulation observed in PWH on ART. In this study, we show that intragenic transcription associated with defective proviruses correlates with induction of IFN type 1 responses in clinical samples, myeloid cell lines, and primary macrophages. Furthermore, RNA-mediated IFN I response is dependent on the MDA5 signaling pathway. We propose a model that innate immune sensing of spurious RNAs from defective HIV-1 proviral genomes perpetuates inflammation in PWH.

## Results

### Cell-associated UTR-deficient transcripts in PWH correlate with increased serum cytokine levels

Our hypothesis is that spurious transcription from defective HIV-1 proviruses induces inflammation. We explored whether there was a correlation between levels of cytokines in PWH and specific subsets of HIV-1 transcripts; those initiated at the LTR and those non-canonical RNAs that were transcribed from intragenic elements. We examined data collected from 22 PWH in a previously described “inflammaging” cohort consisting of PWH on ART for at least 6 months with no detectable viremia (Table 1)[39]. Additional demographic information including, CD4+ T cell count at time of sample collection, and sex assigned at birth are also included in Table 1. As part of the original study, cytokine levels were assessed by ELISA [39]. We focused on those cytokines which had a range of concentrations above the limit of detection, specifically TNFα, IP-10, IL-6, IL-8, and IL-12p70. RNA was isolated from PBMCs for these 22 samples and multiplex-RTddPCR (Fig 1A-B) was used to quantify levels of total HIV RNAs, 5’ untranslated region (UTR)-containing RNA initiating from the LTR, and UTR-deficient RNA arising from alternative promoter-like elements. Univariate analysis for each RNA subset, total RNA, each cytokine, and the demographic factors were run to identify notable correlations (Fig 1C). UTR-deficient RNAs correlated significantly with serum levels of IP-10, TNFα, and IL-8. Total HIV-associated RNA did not correlate strongly with any cytokines, nor did UTR-containing RNA (Fig 1C). Based on these analyses, we ran multiple linear regressions for combinations of HIV-1 RNA populations and these three cytokines, while adjusting for age, biological sex, and time on ART. Age, time on ART, and biological sex did not affect the correlations between UTR-deficient RNAs and TNFα, IP-10, or IL-8 (Sup Tables 5-7). As shown in figure 1, there is a correlation between UTR-deficient RNAs and these cytokines.

**Figure 1:**
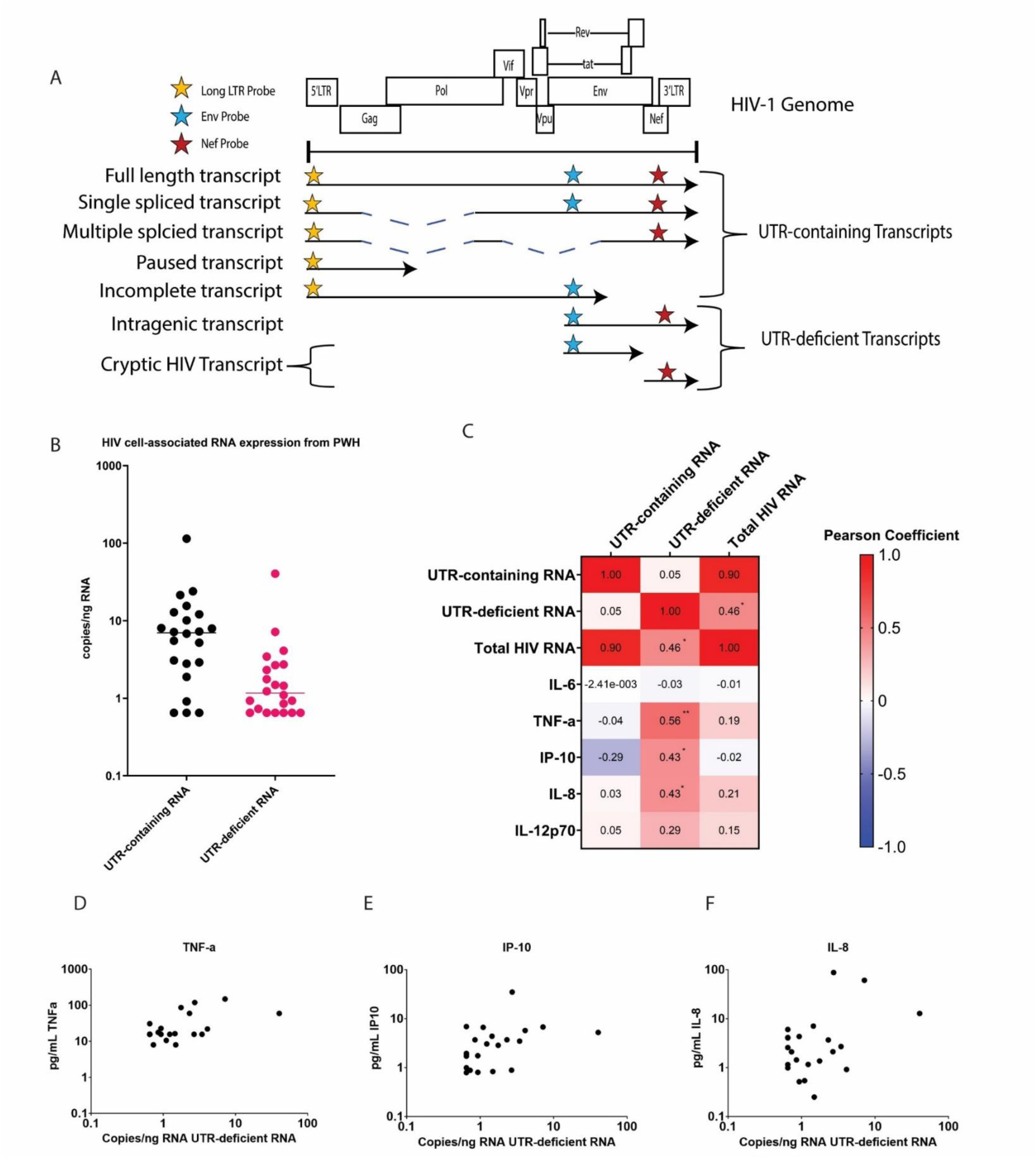
Expression of inflammatory cytokines correlate with UTR-deficient HIV RNA. A) Diagram of the multiplexed RTddPCR assay used to measure HIV-1 transcripts with or without the UTR sequence. Adapted from Kilroy et al. 2021 [26, 61] B) Quantification of HIV UTR-containing and UTR-deficient RNAs from 22 PWH. C) Heat map showing Pearson correlation coefficients between UTR-containing RNA, UTR-deficient RNA, or total HIV-1 RNA modeled against serum cytokine levels. The log of each raw value was taken prior to correlation tests. * represents p-values below 0.05 and ** represents p-values below 0.01 for the univariate analysis. D-F) XY scatter plots of cytokines TNFα (D), IP-10 (E) and IL-8 (F) which were significantly correlated with UTR-deficient RNAs. Untransformed measures of cytokines (pg/mL) and UTR-deficient RNA (copies/ng RNA) are plotted. Axes are on a log-10 scale. Multiple Linear Regression showed significant correlations between UTR-deficient RNAs and IP10 (p = 0.0353, Sup Table 5), TNFα (p = 0.0213 Sup Table 6), and IL-8 (p = 0.0375 Sup Table 7).

**Table 1:** Cohort Description*.

| Characteristic | All PWH (n=22) |
| --- | --- |
| Age years, median (range) | 52 (26-68) |
| Female sex, no. (%) | 8 (36.4) |
| Duration of therapy years, median (range) | 11.49 (2.38-20.63) |
| Absolute CD4 count (cell/mm3), median (range) | 638 (194-1523) |

\*cohort originally described in Pihl and Smith-Mahoney et al. 2024 [39].

### Defective provirus trigger IP-10 production *in vitro*

To explore whether defective HIV-1 proviruses contribute to inflammation, we used CRISPR to generate defective proviruses in infected cells. As proof of concept for our general strategy, we used a Jurkat cell line engineered to constitutively express Cas9. Following infection of these cells, we transfected plasmids that expressed specific gRNAs that targeted the integrated provirus, deleting approximately 1 kilobase of proviral genome from the 5’LTR to the 3’end of *gag* (Fig 2A). The 5’ LTR deletion was confirmed by PCR using primers which flanked the indel sites (Fig 2B) An HIV-1 intact provirus would generate a product of approximately 1.5kb in length, whereas a 400 bp product would be generated from a CRISPR-modified provirus (Fig 2C, Sup Fig 1A-C). We detected a 400bp fragment in infected Jurkats treated with specific guides and observed a decrease in the intensity of the band associated with intact HIV-1 provirus (Fig 2C). Production of interferons from the infected Jurkat cells was assessed by adding spent media from the Jurkat-HIV-1 infected cells to a HEK-293-ISRE-luc cell line which includes an interferon stimulation response element driving a luciferase reporter; addition of type I IFNs to this cell line induces luciferase expression [40]. A modest three-fold induction with supernatants from cells harboring HIV-1 genomes modified by CRISPR was observed. Controls treated with scrambled RNAs as well as cells infected with an HIV-1 clone in which *env* was disrupted by GFP did not induce an interferon response (Fig 2D). These results suggest defective proviruses contribute to a type I IFN response.

**Figure 2:**
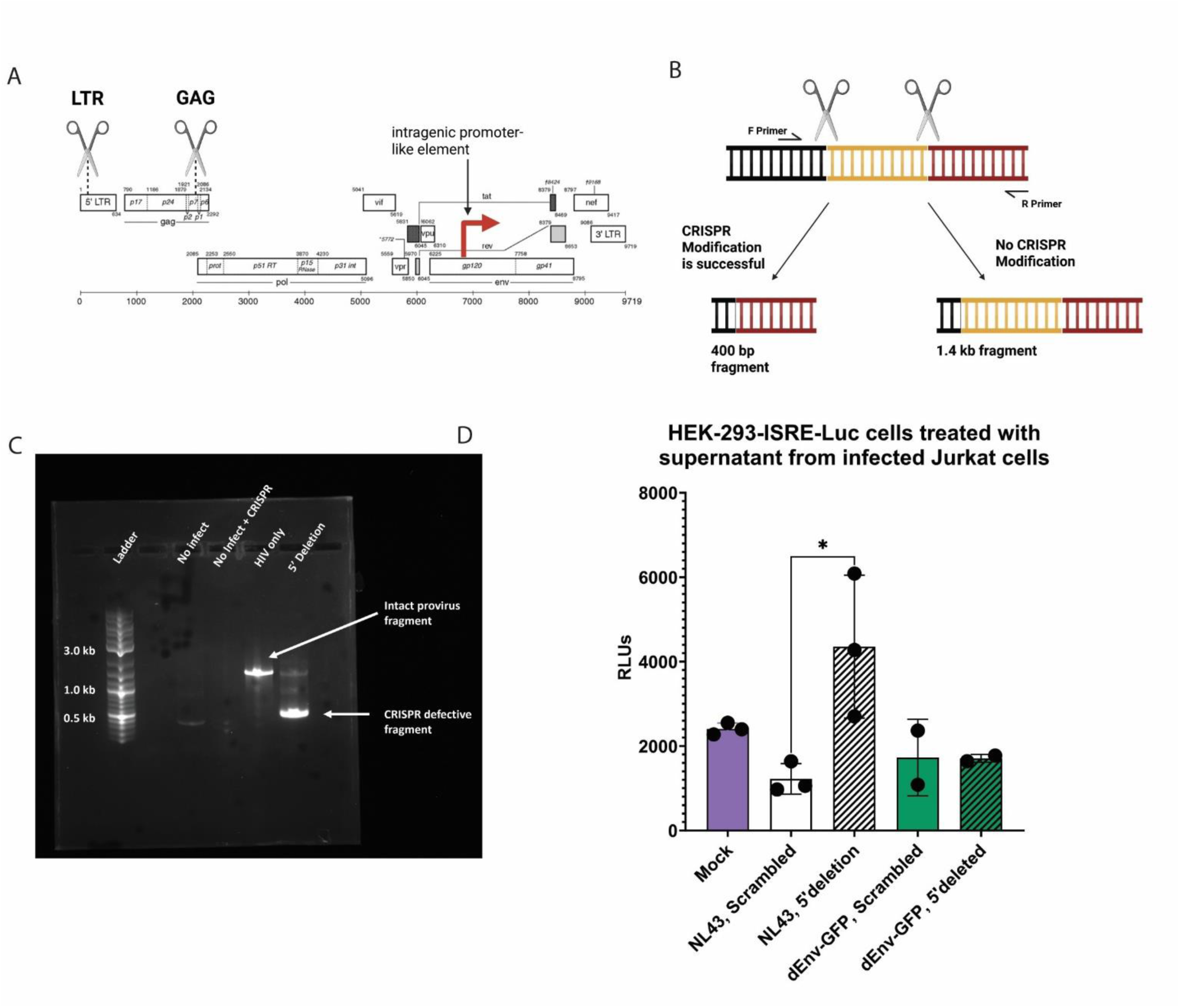
Engineering defective HIV-1 proviruses in Jurkat cells induce type I IFNs. Jurkat-Cas9 cells were infected with VSVg-pseudotyped NL4-3 or VSVg-pseudotyped NL4-3-dEnv-GFP at an MOI of 0.2. CRISPR gRNA-expressing plasmids were nucleofected 24 hours post infection. DNA and supernatants were collected 48 hours post infection. A) Map of the HIV-1 genome annotated with the CRISPR-Cas9 target sites and the intragenic promoter-like element. B) Diagram of the PCR strategy for confirming CRISPR deletion of the HIV-1 genome. C) Agarose gel with PCR products confirming CRISPR modification of HIV-1 provirus in Jurkat cells. D) HEK-293-ISRE-Luc cells were treated with supernatants from Jurkat cells harboring defective proviruses and compared to similarly modified NL4-3-dEnv-GFP virus in which the intragenic element is disrupted. Higher RLUs correspond to higher concentrations of interferons in supernatants. Each data point represents an individual experiment. Error bars represent standard error and ANOVA and paired student’s t tests following a Tukey’s correction. * P < 0.05

HIV-1 associated inflammation and comorbidities has been associated with myeloid cell and macrophage dysfunction including dysregulated production of type I IFNs. To examine whether defective HIV-1 proviruses alter macrophage activity, we utilized CRISPR-Cas9 RNPs to generate MDMs harboring defective proviruses. Generation of defective proviruses were confirmed by PCR as described above and sequencing of that PCR product (Fig A3, Sup Fig 1D-F, Sup Fig 2), which suggested at least a 60% conversion of intact to defective proviruses. HIV DNA levels were determined with an *env* qPCR using a pNL4-3 standard and there was no significant difference in infection between scrambled and 5’deleted conditions (Sup Fig 4A). Induction of IP-10 transcripts and protein were measured by RTqPCR (Fig 3A) and ELISA (Fig 3D), respectively. An increase in IP-10 transcripts and protein was observed when compared to IP-10 expression in unmodified provirus controls and scrambled guide RNA controls. IP-10 transcription increased 75-1,000-fold while IP-10 protein production increased 3-4-fold (Fig 3A-C). IRF1 transcription levels were unchanged between conditions which is expected since this interferon stimulated gene is regulated by post-translational modification (Fig 3E) [41–43]. Controls including scrambled gRNA controls and CRISPR modification to the human gene *hprt* did not induce IP-10 when compared to untransfected cells (Sup Fig 3A-B), with the latter indicating that CRISPR-Cas9 targeting of an endogenous gene does not induce an interferon response. Taken together, these data support the hypothesis that defective HIV-1 proviruses contribute to myeloid cell dysfunction and inflammatory activity.

**Figure 3:**
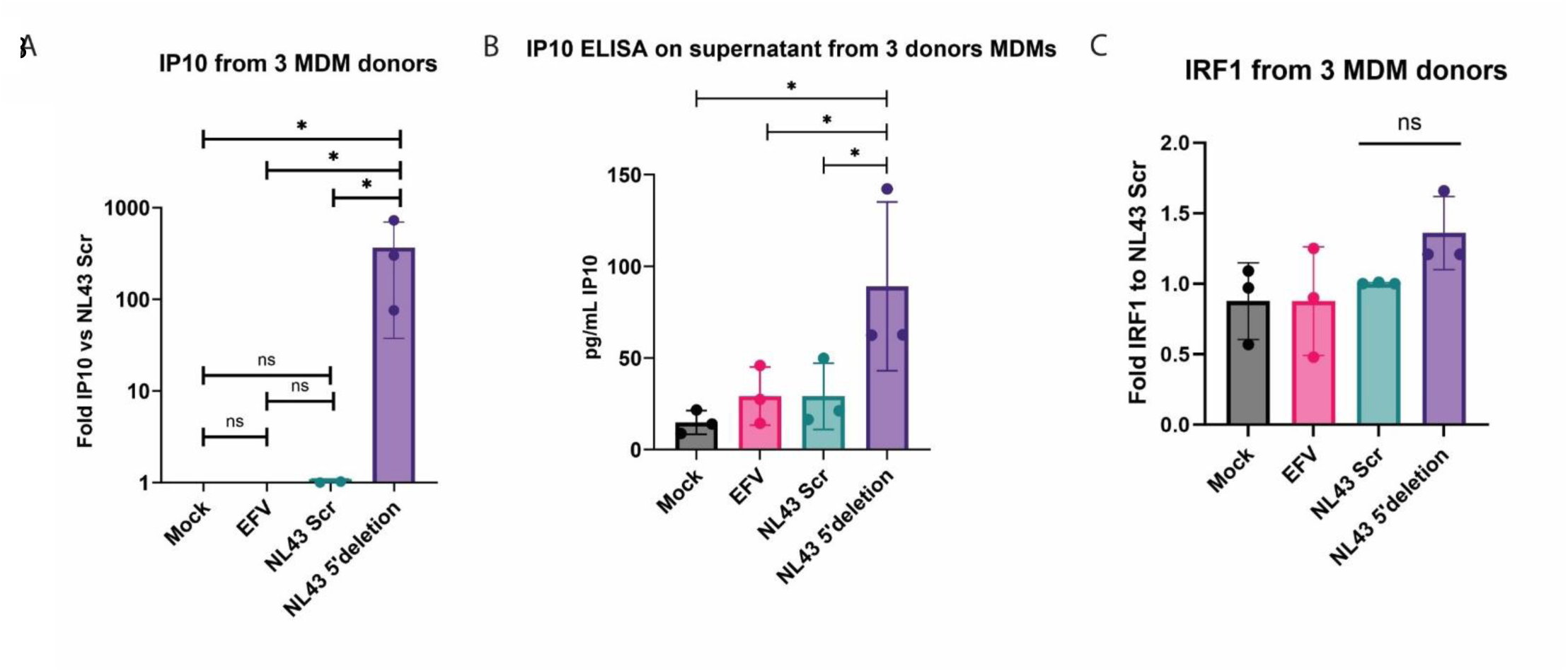
Engineered defective HIV-1 proviruses in MDMs induce IP-10. A) RTqPCR for IP-10. Data is shown as a fold change relative to scrambled gRNA control. Y-axis is a log-10 scale. B) ELISA for IP-10 in supernatant from infected MDMs. C) RTqPCR for IRF1 is shown as fold change relative to scrambled gRNA control. Each data point represents an individual experiment. Error bars represent standard error and ANOVA and paired student’s t tests following a Tukey’s correction. * P < 0.05

### Characterization of UTR-deficient RNA

To characterize the nature of the RNAs generated from defective proviruses, we used bulk RNA from MDM or wild type THP1 cells harboring defective proviruses. To determine if the RNAs contained a 5’7-methylguanosine cap, typically associated with mRNAs, we used recombinant eIF4e protein conjugated to GFP and pulled down 5’capped RNAs using anti-GFP capture beads. Pull down UTR+ and UTR-deficient RNAs were assessed by RTddPCR and compared to bulk RNA from the original population. UTR-deficient RNAs were pulled down by eIF4e in both PMA-differentiated THP1s (Fig 4A) and MDMs (Fig 4B), suggesting that they are 5’capped. The specificity of the pulldown was assessed by comparing levels of leucine-tRNA measured with qPCR in the pre- and post- pulldown RNA populations, since tRNAs do not have a 5’7-methylguanosine cap (Fig 4C-D). We did not detect significant levels of leucine tRNA following 5’cap pulldown although we did detect RPL13a mRNA, a housekeeping gene that is 5’ capped (Sup Fig 4D-E). To determine whether these RNAs are polyadenylated, we generated cDNA using either polydT primers or a combination of random hexamers and polydT primers and compared the ability to amplify UTR-containing and UTR-deficient HIV-1 RNAs using multiplex RTddPCR. We observed intragenic-derived RNAs in both samples (Fig 4E-F), consistent with the non-canonical RNAs having a 3’polyA tail, as observed in our previous study [38]. Overall, these data indicate that intragenic-derived transcripts are modified with a 5’cap and polyA tail.

**Figure 4:**
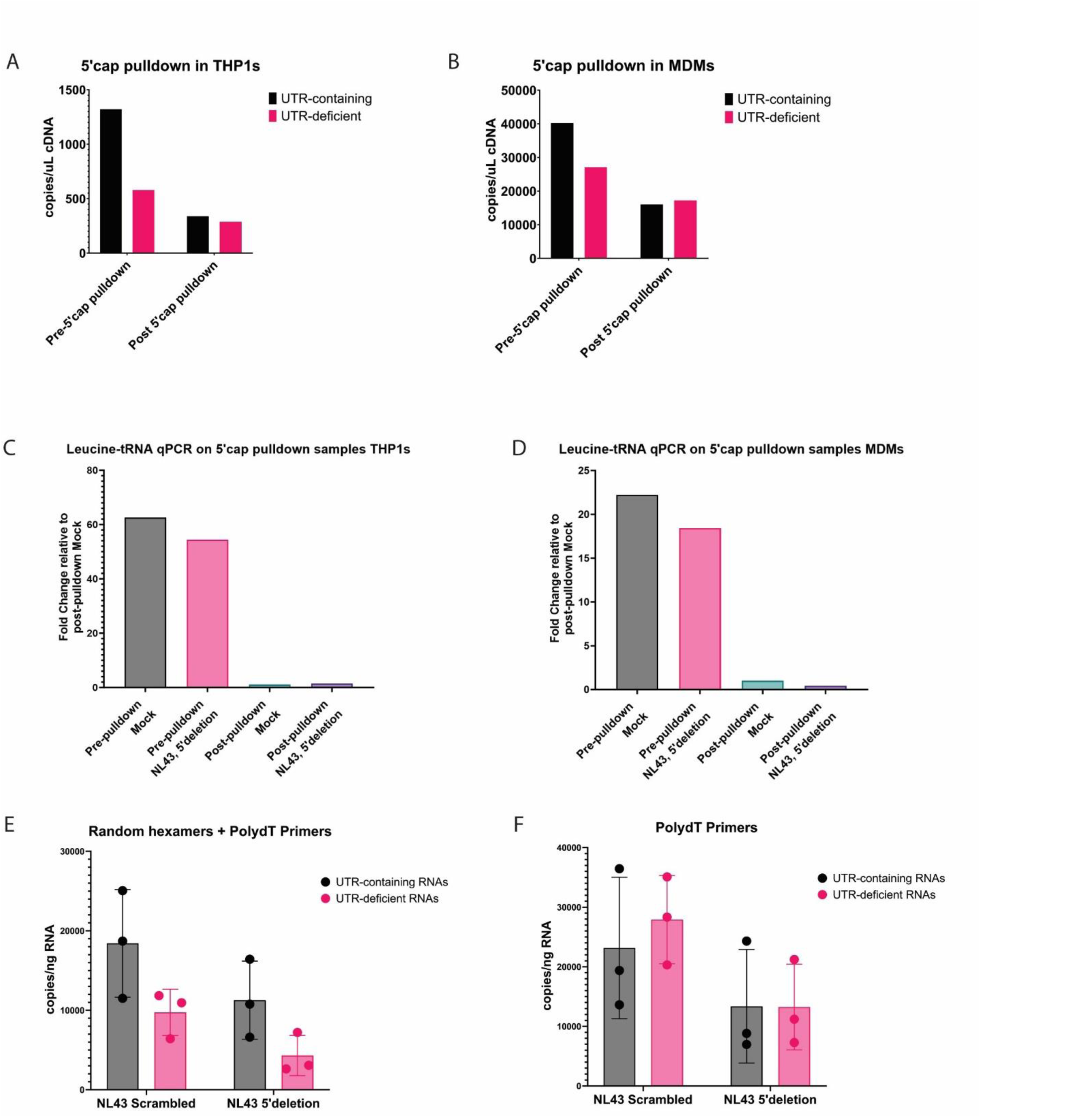
Defective HIV-1 RNAs are capped and poly-A tailed. A & B) RNA was isolated from PMA-differentiated THP1 cells (A) or MDMs (B) harboring CRISPR-modified defective provirus. A portion of this RNA was bound to eIF4e-GFP coated beads which were used to capture 5’capped mRNA. Levels of UTR-containing and UTR-deficient transcripts were determined using multiplex RTddPCR. Results are shown for RNAs pre- and post-pulldown. C & D) RTqPCR for Leucine-tRNA which does not contain a 5’cap in PMA-differentiated THP1s (C) or MDMs (D) pre- and post-pulldown for 5’capped RNAs. E-F) RNA was isolated from MDMs infected with NL4-3 and transfected with scrambled or 5’deleting CRISPR RNPs. cDNA was prepared using a mixture of random hexamers and polydT primers (E) or polydT primers alone (F), then analyzed with multiplexed RTddPCR.

### MDA5 and MAVS are necessary for innate immune responses to defective HIV-1 proviruses

RNAs can act as PAMPs that are recognized by intracellular PPRs. For example, MDA5 has been shown to recognize double stranded RNAs. Furthermore, the MDA5-MAVS signaling cascade has been shown to be necessary for detection and induction of type I IFN responses to HIV-1 intron containing RNAs. To determine if the MDA5/MAVS pathway was similarly necessary for the type I IFN response triggered by defective HIV-1 proviruses, we utilized THP1 cell lines that had either MDA5 or MAVS knocked down [37]. THP-1 cells were infected with HIV-1 and treated with RNPs with Cas9 and guide RNAs as described above. Knockdown was confirmed with PCR (Sup Fig 1G-I). Controls included cells treated with scrambled guides and cells not treated with CRISPR or infected with HIV-1. Compared to wild type THP1s, cells lacking MDA5 or MAVS were significantly compromised in their ability to induce IP-10 transcription in response to defective HIV-1 proviruses (Fig 5 A-C). Overall infection was not affected by CRISPR modification as shown by a qPCR quantifying copies of *env* (Sup Fig 4C-D). These data support the hypothesis that intragenic-derived RNAs are sensed by MDA5 which signals through MAVS to trigger a type I IFN response. Our results are consistent with previous studies showing that MDA5 is necessary for detection of HIV-1 RNA mediated induction of IFNs [30, 31, 44].

**Figure 5:**
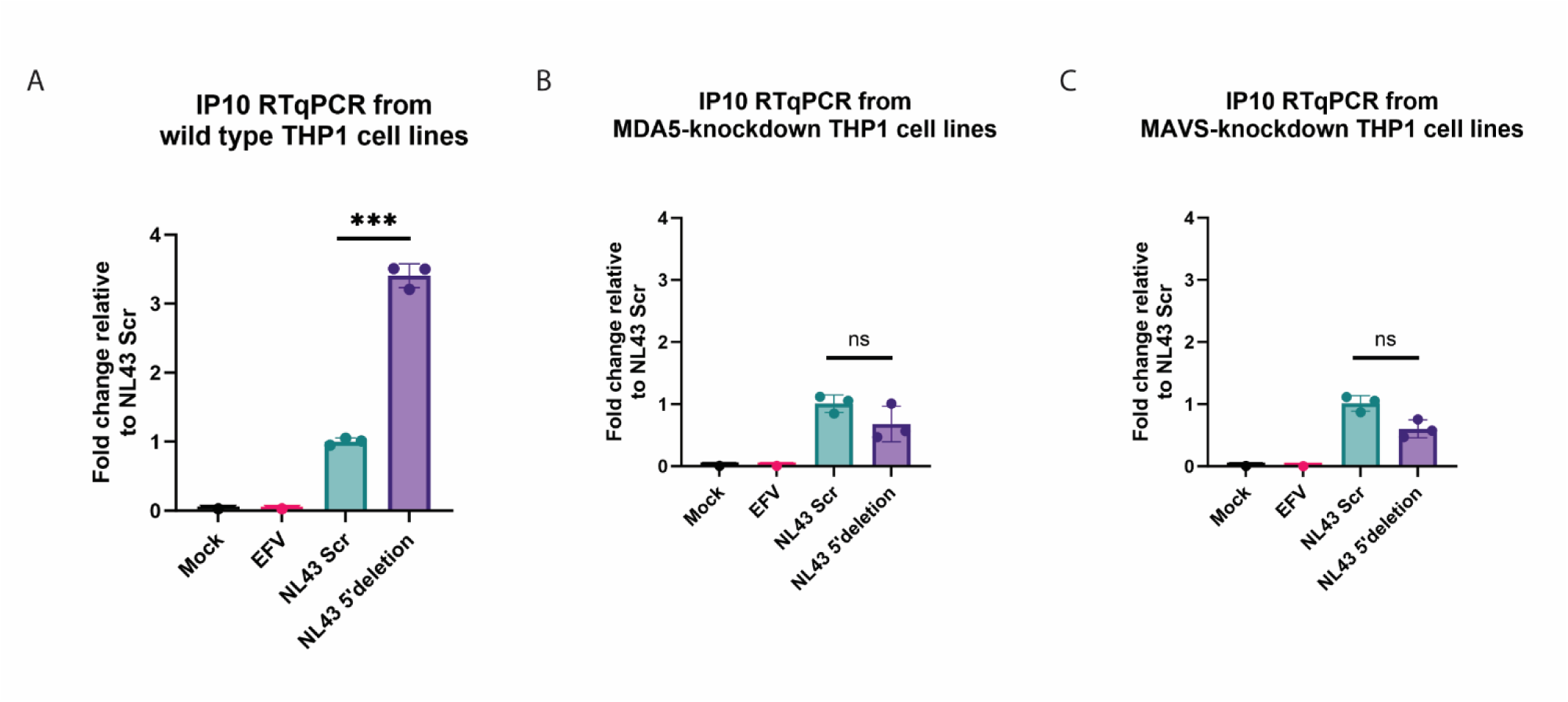
MAVS and MDA5 are necessary for induction of IP-10 by defective HIV-1 proviruses. IP-10 RTqPCR using RNA from A) wild type PMA-differentiated THP1s, B) MDA5 knockdowns, and C) MAVS knockdowns, infected with NL4-3 and transfected with either scrambled CRISPR RNPs or 5’deleting CRISPR RNPs. Each data point represents an individual experiment. Error bars represent standard error and ANOVA and paired student’s t tests following a Tukey’s correction. *** P < 0.001

## Discussion

PWH on suppressive ART experience a wide range of comorbidities posited to be caused by chronic low-grade inflammation [17, 20, 36, 45]. Previously, our group demonstrated that *env* had an intragenic cis-element that functioned as a transcriptional promoter and could facilitate production of non-canonical HIV-1 transcripts [38]. To explore whether these RNAs had a functional consequence in infected cells, we utilized CRISPR-Cas9 to delete the 5’ LTR of HIV-1 proviruses. Cells harboring these defective HIV-1 proviruses triggered increased expression of IP-10. These findings were observed in various cell types, including macrophages, which have been implicated in several end-organ chronic diseases associated with HIV-1 infection and aging [18, 19]. Knocking down MDA5 or MAVS abrogated these responses, indicating that sensing of UTR-deficient RNAs is mediated via the MDA5-MAVS signaling cascade, a pathway implicated in sensing partially spliced HIV-1 transcripts. Consistent with a role in perpetuating HIV-1 associated comorbidities, we observed a correlation between non-canonical UTR-deficient RNAs and elevated levels of IP-10, TNFα, and IL-8 in samples from PWH receiving ART.

Several groups have suggested that innate immune sensing of viral RNA, particularly unspliced HIV-1 RNA, mediates chronic inflammation. Sensing of exported HIV-1 unspliced RNA in DCs and MDMs by MDA5 and MAVS drives production of pro-inflammatory interferon stimulated genes including IP-10, ISG15, and MX1 [10, 30, 31, 37, 46]. Total HIV-1 RNA has been shown to correlate significantly with levels of proinflammatory cytokines in serum from PWH in some studies, although others show no correlations with total HIV-1 RNAs [47–50]. While total HIV-1 RNAs did not reach significant correlations in this study, there was a trend of positive correlation between total HIV RNA and TNFα. However, prior studies have not included UTR-deficient RNAs as a key predictor of cytokine levels, instead using the UTR sequence to identify HIV-1 RNAs or a single target site within the provirus [48, 49, 51]. Furthermore, the structure of the 5’ leader region of viral RNA plays an important role in triggering the MDA5 response [44]. Unspliced RNAs and active HIV-1 protease also trigger inflammasomes, interleukin 1 beta (IL-1β) production and pyroptosis [28, 29, 44, 52, 53]. We propose that HIV-1 mRNAs through multiple mechanisms activate and dysregulate multiple intracellular innate immune responses in myeloid cells such as DCs and macrophages, to synergistically drive inflammation.

Although our data support that HIV-1 noncanonical RNA is inducing an interferon response, it is possible that these mRNAs are translated and that these proteins contribute to HIV-1 pathogenesis. The mRNAs generated from the intragenic promoter with *env*, contain an intact open reading frame (ORF) and are 5’ capped and polyadenylated, therefore, having features of mRNAs that are translated. We previously described the detection of proteins associated with these *env* transcripts by immunoblotting using anti-sera from PWH [38]. Several studies have shown that defective proviruses have the capacity to generate peptides which are presented as antigen, eliciting CD8+ T cell responses [14, 26, 27, 54, 55]. Several ORFs generated from alternative splicing and specific defects give rise to full length HIV-1 proteins, truncated HIV proteins, and cryptic HIV-1 peptides, and these could shape the innate immune response and the anti-HIV-1 T-cell responses [14, 56, 57]. For example, Gag and Nef proteins have been detected in PWH receiving suppressive ART and are suggested to trigger innate immune responses contributing to a failure of CD4+ T cells in PWH who are immunological non-responders [14, 58]. Furthermore, Pollock et al suggested that class I presentation of peptides from defective proviruses elicit CTL responses either diverting or perpetuating CD8+ responses in PWH [54]. We speculate that a subset of RNAs we observed, in addition to triggering a response through MAVS, are translated into proteins that directly or indirectly skew innate and T cell responses.

Based on our findings, we propose a model for defective provirus-driven inflammation. An intragenic element within *env* remains transcriptionally active, even during latency and suppressive ART regimens. RNAs transcribed from this element are modified with a 5’cap. MDA5 senses these RNAs, triggering an interferon response through MAVS, leading to increased production of interferons and IP-10. In addition, we hypothesize that these RNAs are translated and further skew innate and adaptive responses. The continuous expression from this region in PWH drives chronic inflammation. Overall, we propose that these HIV-1 RNAs contribute to an inflammatory microenvironment that directly perpetuates HIV-1 associated comorbidities. These studies indicate that persistent defective HIV-1 proviruses are critical in long-term chronic HIV-1 pathogenesis and their role in HIV-1 associated comorbidities need to be considered as ongoing treatments and cure strategies are explored (Figure 6).

**Figure 6:**
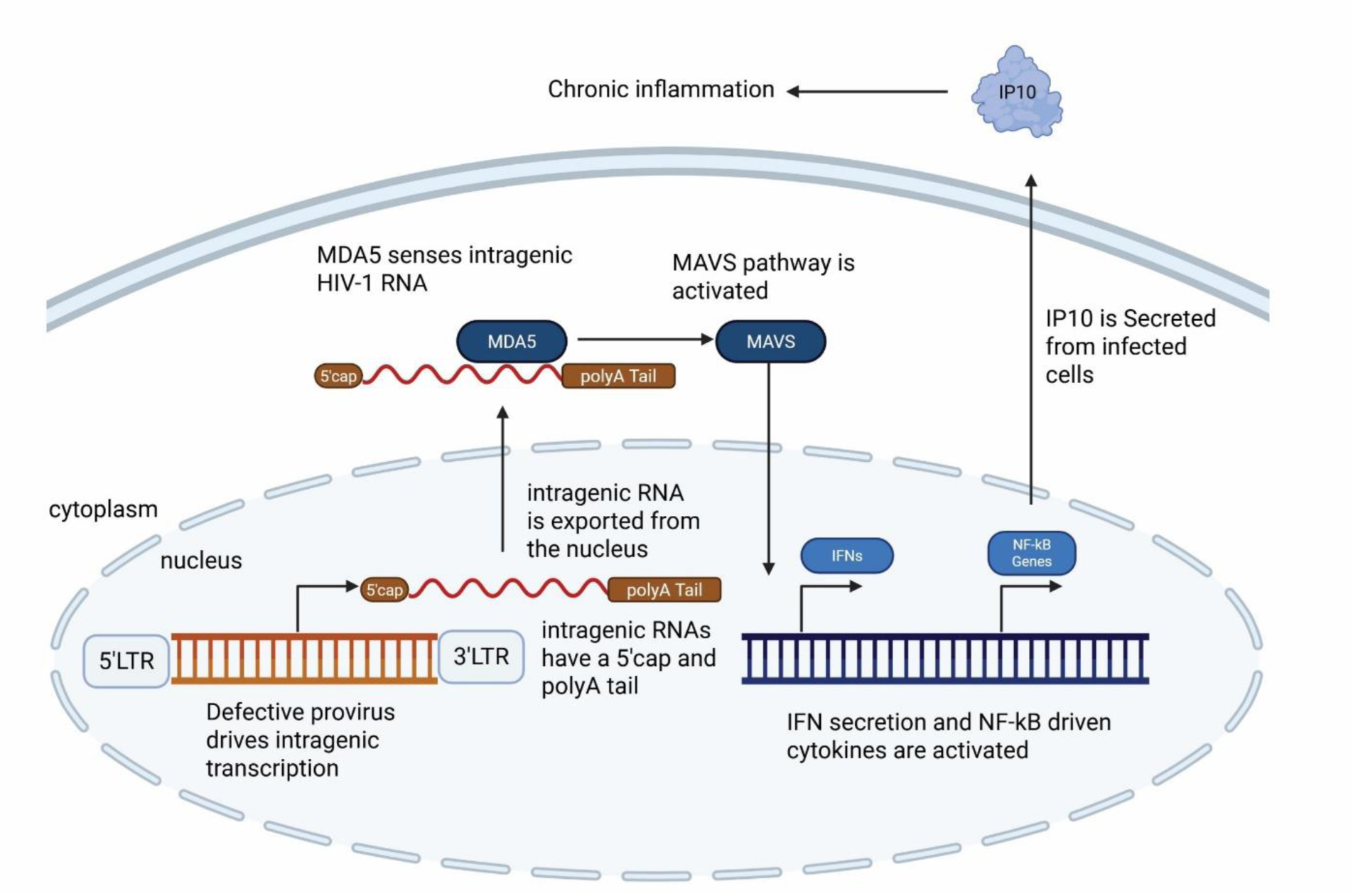
Proposed model for defective provirus-driven inflammation.

## Materials and Methods

### Virus preparation

HIV-1 viruses were packaged and pseudotyped with VSVg envelope by transfecting HEK-293T cells as previously described [38]. Viruses were concentrated through a 20% sucrose gradient. Virus titers were obtained as previously described by infecting a CEM-R5-GFP reporter line which produces GFP in response to Tat production and measuring infection percentage by flow cytometry to determine the multiplicity of infection (MOI) [38]. All infections were performed with an MOI of 1.0 except for Jurkat cells, which were infected with an MOI of 0.2.

### Infection with HIV-1

Virus was added to Jurkat cells and cultures spinoculated at 1200xg for 90 minutes. Cells were washed with PBS and resuspended in fresh media following spin. Samples were harvested for supernatants, RNA, and DNA at the indicated times Differentiated THP1 cell lines and MDMs were incubated with virus overnight. Cells were washed with PBS and provided fresh culture media. CRISPR-Cas9 RNPs were added 24 hours post infection. THP1 samples were harvested 48 hours post infection. MDM samples were harvested 72 hours post infection.

### Isolation of PBMCs

Blood packs were obtained from Rhode Island Blood Center. Blood was diluted in PBS and separated over a Ficoll-Hypaque gradient (Stem Cell Technologies Catalog No. 7801). PBMC buffy coats were removed from the interface of plasma and the density gradient and washed with PBS. Cells were treated with red blood cell lysis buffer (155mM NH_2_Cl, 12mM NaHCO_3_, 0.1mM EDTA), washed with PBS, and resuspended in selection media (PBS +2% FBS +2mM EDTA). PBMCs were stored in liquid nitrogen for long term storage. For the purification of CD14+ monocytes a Stem Cell CD14+ human selection kit was used (Stem Cell Technologies Catalog No. 17858). Monocytes were plated in RPMI with 1% Pen/Strep (Thermo Fisher catalog no. 15140122), 1% L-glutamine (Thermo Fisher catalog no. 25030081), 5% heat-inactivated FBS (Phoenix Scientific catalog no. PS-100), 10% human AB serum (Sigma Aldrich catalog no. H4522), and 50ng/mL M-CSF (Biolegend catalog no. 574804) for 6-7 days to differentiate into MDMs. MDMs were maintained on complete RPMI (10% FBS, 1% Pen/Strep, 1% L-glutamine).

### Cell lines

THP1 (ARP-9942) cell lines were maintained in RPMI with 1% Penicillin/Streptomycin (Invitrogen), 1% L-glutamine, and 10% FBS. Polyclonal THP1-cell lines constitutively knocked down for MAVS or MDA5 were engineered as previously described [37]. Prior to infection, THP1 cell lines were differentiated overnight using 100nM PMA before reseeding into 12-well plates for infection.

A polyclonal Jurkat cell line stably expressing Cas9 was generated by infecting Jurkat cells (ATCC TIB-152) with lentivirus containing the pLX_311-Cas9 vector (Addgene 96924). For lentiviral packaging, HEK293T cells (ATCC CRL-11268) were grown in DMEM + 10% FBS + 2 mM GlutaMAX™ Supplement and transfected with a DNA mixture containing the pLX_311-Cas9 vector, the packaging plasmid psPAX2 (Addgene 12260), and envelope plasmid VSV.G (Addgene 14888) using polyethyenimine. The resulting lentivirus was concentrated using the Lenti-X™ Concentrator reagent (Takara 631232) according to the manufacturer’s protocol. Jurkat cells were transduced using 1:5, 1:10, 1:50, 1:100, and 1:500 dilutions of Cas9 lentivirus to RPMI 1640 + FBS + GlutaMAX™ containing 10 µg/mL polybrene and incubated for 48 hours at 37°C, 5% CO_2_. Cells were expanded for several days before selecting for pLX_311-Cas9 vector in RPMI 1640 + FBS + GlutaMAX™ containing 10 µg/mL blasticidin. Media was changed every 3 days until cells in the non-transduced control was 0% viable, proceeding with the cell population with the highest viability. All cells were cultured at 37⁰C, 5% CO_2_.

### HEK-293-ISRE-Luc bioassay for interferons

HEK-293 cells were engineered with a lentivirus construct that contains an ISRE upstream of a luciferase gene as described in Larocque et al 2011 [40]. Supernatants are incubated with these cells for 24 hours in a white-walled 96-well plate. Media/supernatant was aspirated from these plates and 30 µL of BrightGlo reagent (Promega Catalog No. E2610) were added to lyse the cells and measure luciferase activity. A standard curve was generated by adding a serial dilution of recombinant interferon alpha to the reporter line.

### 5’ cap pulldown assay

5’7meG capped mRNA was isolated using a modification of methodology from Blower et al [59]. 500 ng of RNA samples were denatured by heating to 70^0^C for 10 minutes then placing on ice. 250 μL of Buffer A (10 mM potassium phosphate buffer pH 8.0, 100 mM KCl, 2 mM EDTA, 5% glycerol, 0.005% Triton X-100, and 1.3% poly(vinyl) alcohol 98–99% hydrolyzed), 1 μL of RNase Inhibitor, and 2 μg of GFP-tagged high-affinity 5′ cap–binding eIF4E (K119A) was added. Following incubation on ice for 30 minutes, 12.5 μL of ChromoTek GFP-Trap® Magnetic Particles M-270 was added. Following another 30-minute incubation on ice, beads were washed 5 times with 1 mL of ice-cold Buffer A using a magnet to enrich for capped mRNAs. The resulting pellet was resuspended in Trizol, and RNA was purified with Trizol extraction. cDNA was prepared and RTddPCR was used to quantify HIV-associated RNA. Additionally, an RTqPCR assays using Leucine-tRNA and RPL13a primers for non-specific and specific controls were ran to verify the specificity of binding of capped RNAs.

### Preparation of cells for nucleic acid isolation

Following infection with HIV-1, cells were treated with 1 U/mL DNAse and 100 μM MgCl_2_ for at least 30 minutes prior to harvesting. Adherent cells were pelleted at 500xg for 5 minutes. Supernatant was removed from adherent cells and centrifuged at 500xg for 5 minutes to pellet cell debris. Supernatants were then transferred to fresh tubes and stored at −20°C. RNA and DNA were isolated using the Qiagen AllPrep DNA/RNA Kit (Catalog Number 8020wereor Zymo Quick-DNA/RNA Miniprep kit (Zymo, Catalog number D7001) following manufacturer protocols. For some experiments, RNA was isolated using the Maxwell RSC simplyRNA Cells kit (Promega, catalog number: AS1390) and DNA was isolated using the Maxwell RSC Genomic DNA kit (Promega, catalog number: AS1880) according to manufacturer protocol.

### CRISPR-Cas9 modification of integrated HIV

Jurkat-Cas9 cells were nucleofected using the Amaxa Lonza Nucleofector II with plasmids that expressed CRISPR gRNAs in a BW1721 backbone. These plasmids were generated by digesting BW1721 with BbsI, which cuts two sites between a U6 promoter and the gRNA scaffold sequence. Primers were designed with the target sequence for the gRNA and overhangs complementary to the BbsI cut sites (Sup Table 1-2). These primers were incubated with T4 PNK in T4 ligase buffer using T4 PNK at 37°C for one hour. 4µL of 0.5M NaCl was added to the solution, and the mixture was heated to 95°C for 5 minutes. The temperature was decreased by 5°C every 2 minutes until reaching room temperature to mediate primer annealing. SDS was used to denature the enzyme, then PCR cleanup performed to purify the annealed primers using Monarch Spin PCR and DNA Cleanup Kit (New England Biolabs Catalog No. T1130L). This product and the BW1721 backbone were digested separately with BbsI at 37°C for 2 hours then purified via PCR cleanup kit (New England Biolabs Catalog No. T1130L). The annealed primer insert and digested plasmid backbone were ligated overnight at room temperature using T4 DNA Ligase. Plasmids were validated by Sanger sequencing using a U6 promoter-specific primer to initiate sequence. Plasmids were transformed into *E. coli* and prepared from large cultures of the transformed bacteria. Plasmids were nucleofected into Jurkat-Cas9 cells per manufacturer protocols.

For THP-1 and MDMs, approximately one million cells were plated per well of a 12-well plate. Cas9, gRNAs, and Cas9 Plus Enhancer were mixed and delivered to cells as a RNP by CRISPRMAX lipofectamine (Thermo Fisher Catalog No. CMAX00003). Full length gRNA sequences are in Supplemental Table 2. Supplemental Table 3 shows ratios of Cas9 and gRNAs.

Modification of HIV proviruses by CRISPR was confirmed by PCR using primers which flank the CRISPR-Cas9 cleavage site. 10-500ng of genomic DNA and 1.25µL of each primer was added to 25µL of Q5 2X master mix (New England Biolabs, catalog number M0492S) and the total volume adjusted to 50µL. Samples were amplified for 30-45 cycles with an annealing temperature of 61°C. PCR-amplified DNA was loaded onto a 1% agarose gel and ran at 80 volts to visualize PCR products. Fragments were also confirmed by sequencing the PCR product.

### Sample Analysis

Multiplexed RTddPCR assay for the quantification of HIV-1 expression and splice variants was performed as previously described [60]. The assay is outlined in Figure 1A.

RTqPCR was performed using SYBR Green Master Mix (AB Clonal Catalog No. RK21203).

Primers used for the PCRs are shown in Supplemental Table 1.

### ELISA Measurement of Cohort Serum Cytokines

ELISA data for donor serum cytokine was previously published data by Pihl and Smith-Mahoney et al. [39].

### ELISA Measurement of IP-10 from *in vitro* samples

The DuoSet Solid Phase Sandwich ELISA kit for CXCL10/IP-10 (R&D Systems, Catalog no. DY266) was used to quantify levels of IP-10 protein in the supernatant from *in vitro* experiments per manufacturer protocols.

### Statistical Analysis

All statistics were calculated using GraphPad Prism software. PCR and ELISA data were processed using ANOVA and paired student’s t tests following a Tukey’s correction. Analysis for clinical data correlations was run using Pearson correlations and multiple linear regression.

Cytokine values and RNA values which were below the limit of detection were set to the lowest measurable value for each assay. The logarithm of concentrations of cytokines in serum (pg/mL) and copies/µL/ng of HIV-1 cell-associated RNA were taken for analysis. These factors were tested for normality using D’Agostino and Pearson test, Anderson-Darling test, Shapiro-Wilk test, and Kolmogorov-Smimov test (Sup Table 4). The cytokines and RNA copies were normally distributed, although UTR-deficient RNAs, IL-12, and age at time of enrollment were not normally distributed (Sup Table 5). Univariate analysis was run using a Pearson Correlation test. Multiple Linear Regression models were run for each cytokine to determine if each subset of HIV-1 cell-associated RNA and demographic factor were good predictors.

## Acknowledgements

We thank Drs. Rachel Yuen and Alex Olson for their assistance in processing and consolidating data from clinical samples. We also acknowledge Brian Tilton at the BUMC Flow Cytometry Core Facility and the assistance provided by Providence/Boston CFAR Scientific Core (P30 AI042853). This project was funded in part by Boston University’s Undergraduate Research Opportunities Program to AD and National Institutes of Health grants R01 AI187175 and DA055488 to AJH & SG.

---

**Supplemental Figure 1:**
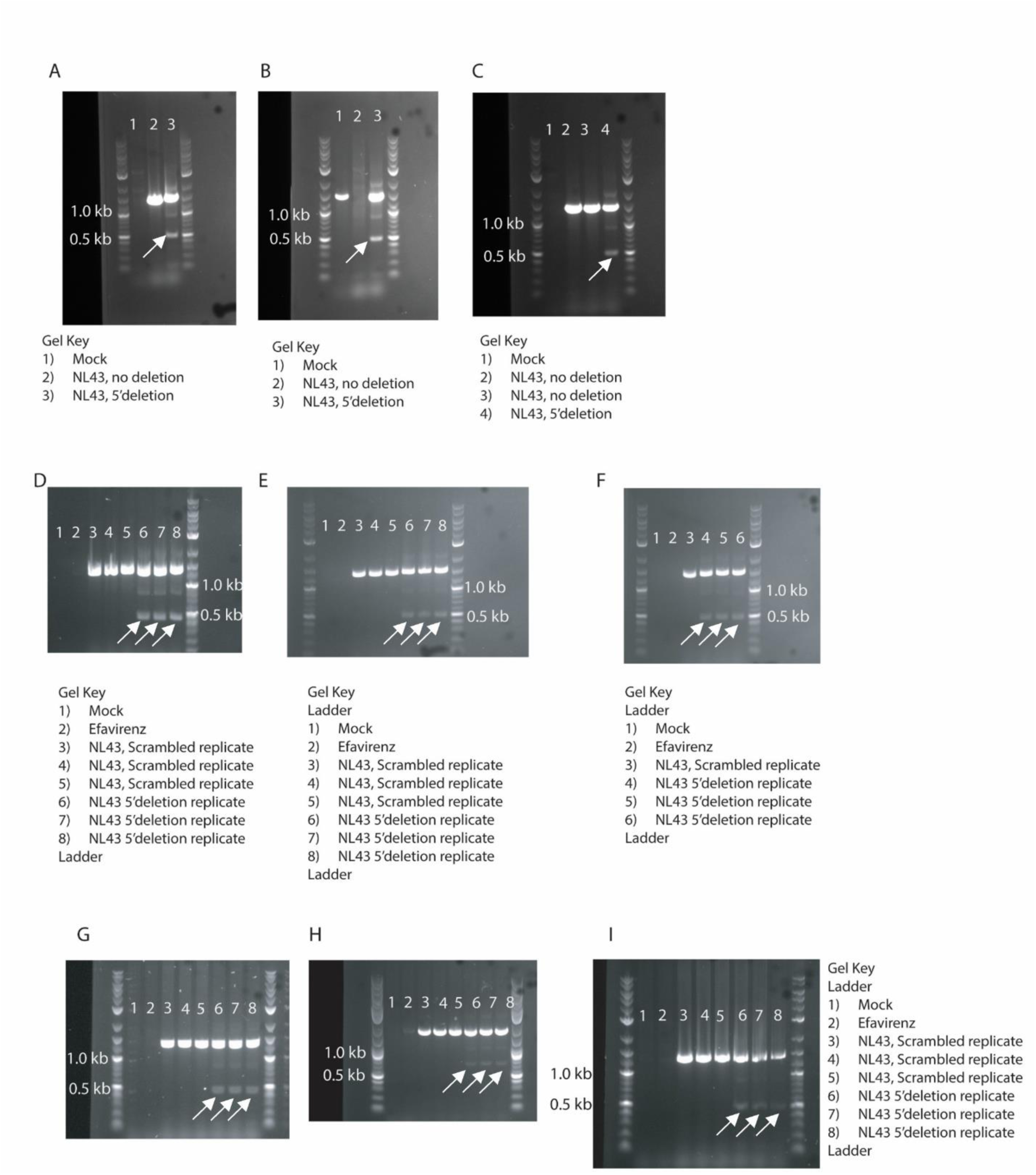
Confirmation of knockdowns by PCR. White arrows indicate PCR amplicons arising from CRISPR-modified HIV-1 proviruses. A-C) PCR was used to amplify genomic DNA as shown in Fig 2B from Jurkat-Cas9 cells infected with NL4-3 and transfected with either scrambled or 5’deleting gRNA-expressing plasmids. Each panel represents a different replicate within Jurkat-Cas9 cells. D-F) PCR products from genomic DNA isolated from MDMs harboring full length or modified HIV-1 provirus. Each gel represents a unique MDM donor with 3 replicates per infected condition per donor. G-I) PCR products from genomic DNA isolated from PMA-differentiated THP1s (G), PMA-differentiated MAVS-knockdown THP1s (H) and PMA-differentiated MDA5-knockdown THP1s (I).

**Supplemental Figure 2:**
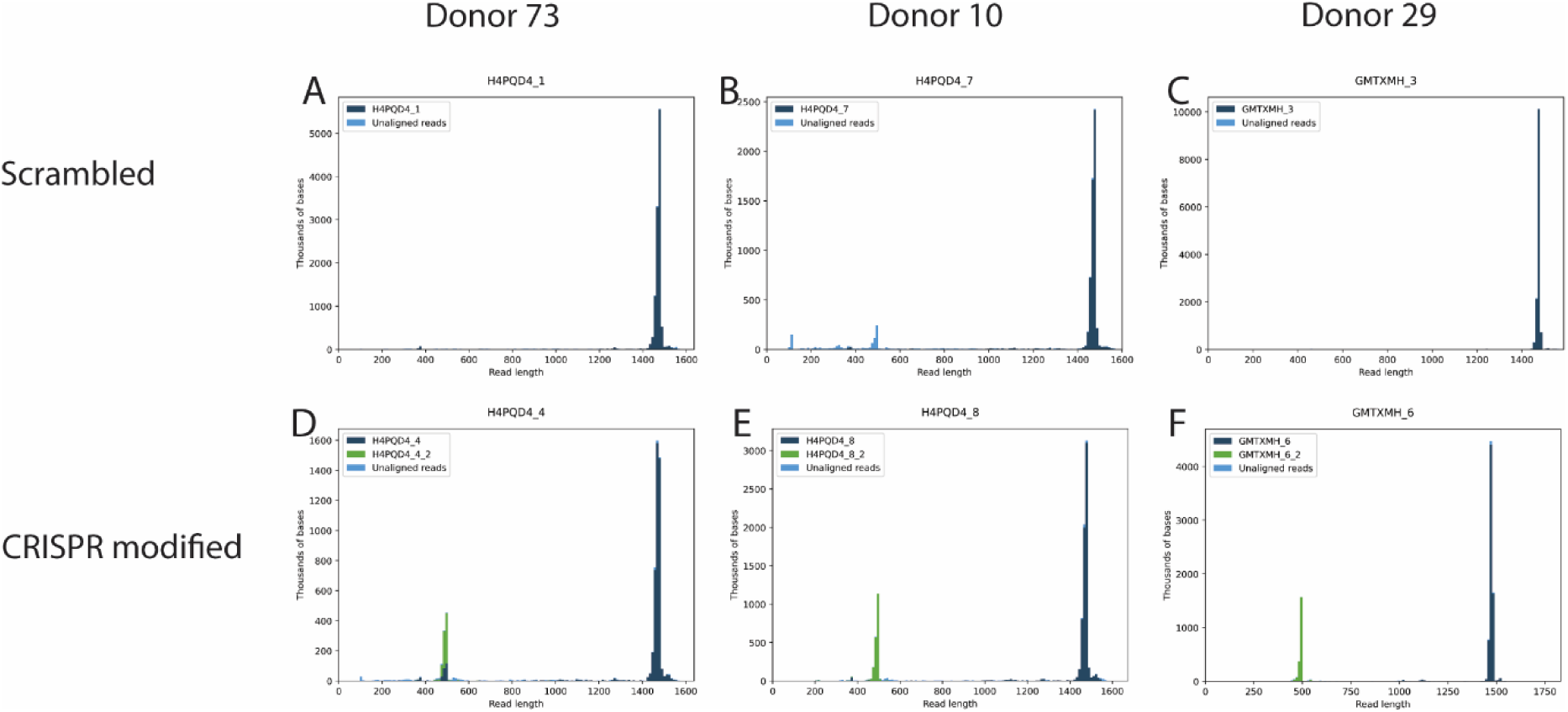
Confirmation of modified HIV-1 proviruses by sequencing. Histograms showing the distribution of PCR amplicon reads from MDM genomic DNA as described in the methods. A-C) represents samples from a single donor infected with NL4-3 and transfected with scrambled CRISPR-Cas9 RNPs. D-F) represents samples from a single donor infected with NL4-3 and transfected with 5’deleting CRISPR-Cas9 RNPs.

**Supplemental Figure 3:**
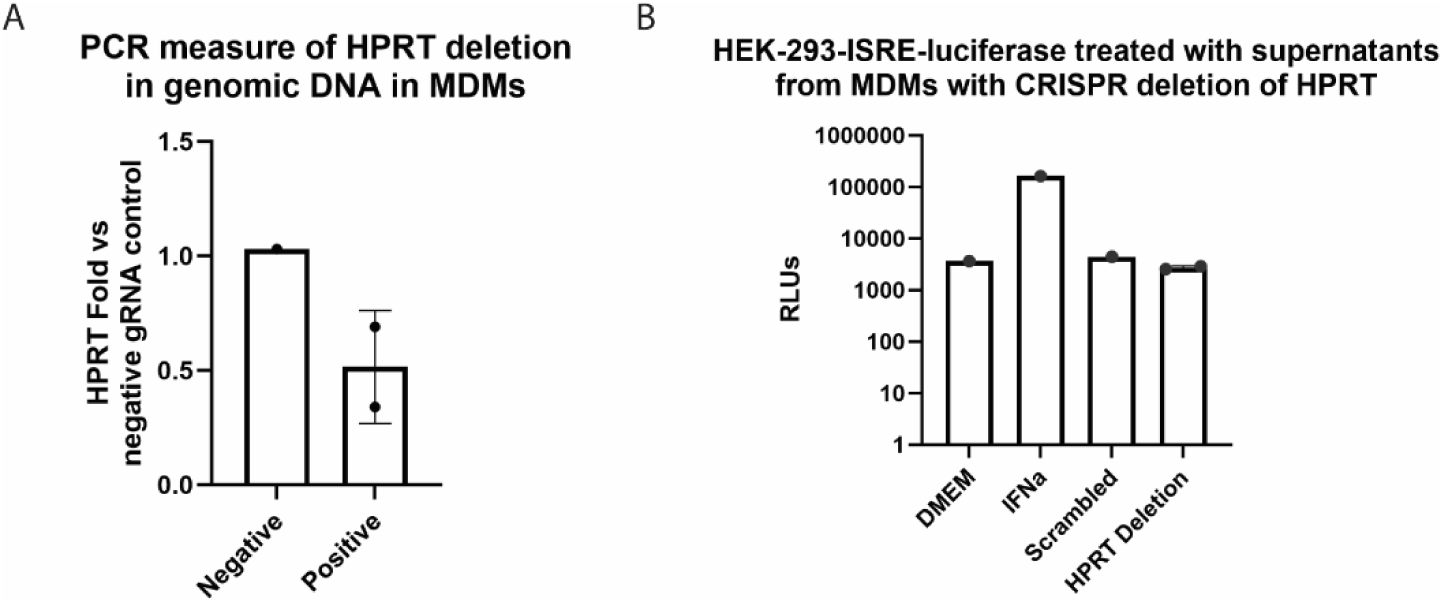
Targeting *hprt* in MDMs does not induce interferons. A) qPCR confirming knockdown of endogenous human gene *hprt* from genomic DNA in MDMs. *hprt*-targeting CRISPR-Cas9 RNPs were formed as described in the methods. DNA was harvested 48 hours post transfection. B) Supernatant from modified MDMs was added to HEK-293-ISRE-Luc cells and compared to the addition of DMEM (negative control) and recombinant Interferon alpha (positive control) to the reporter cell line.

**Supplemental Fig 4.**
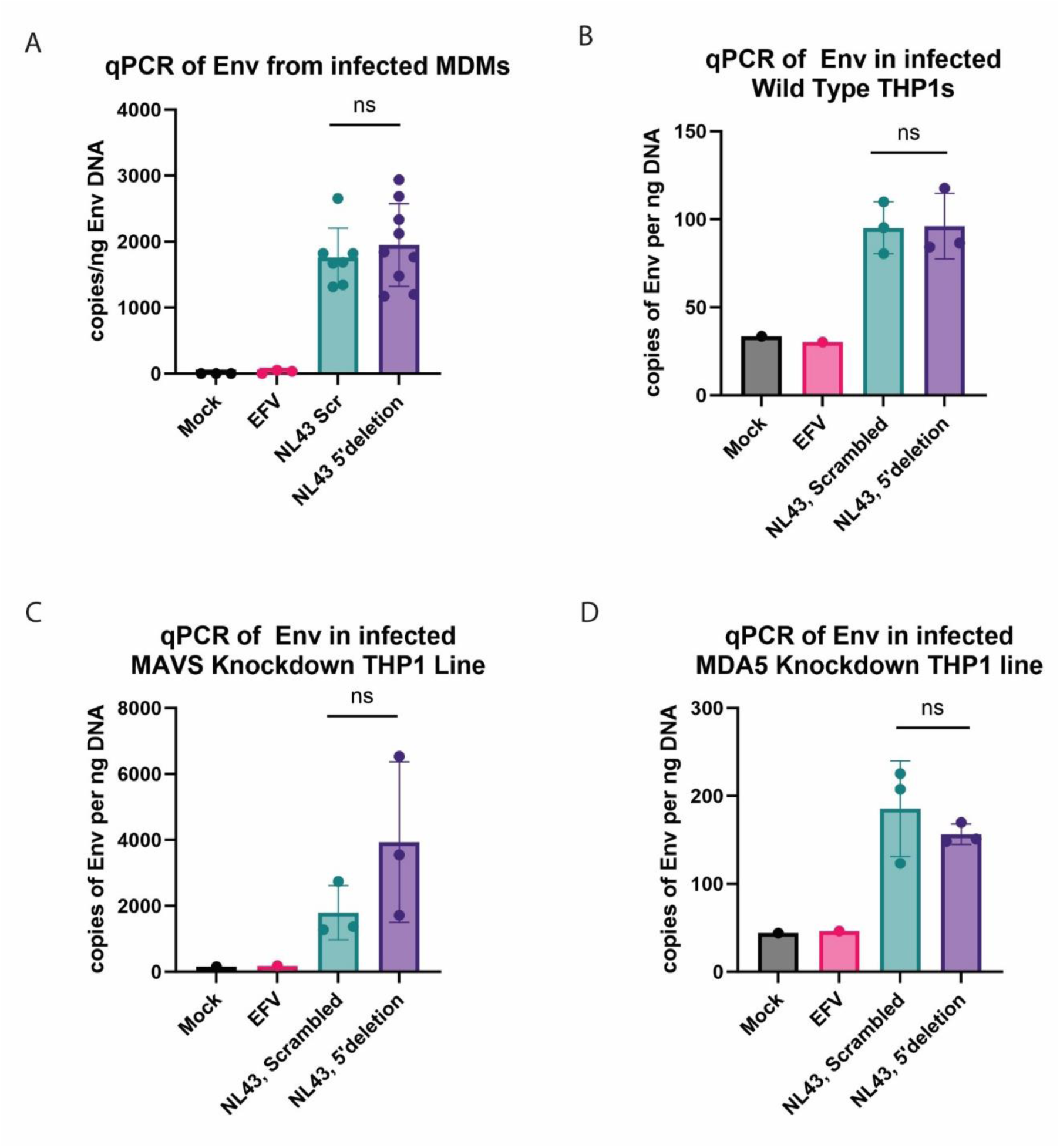
CRISPR RNP treatment did not impact HIV-1 infection. qPCR was used to quantify copies of *Env* DNA per ng input genomic DNA to determine the overall copies of provirus following CRISPR modification. Titration of NL-43 was used determine *env* copy. Data are from MDMs (A), PMA-differentiated THP1s (B), PMA-differentiated MAVS-knockdown THP1s (C), and PMA-differentiated MDA5-knockdown THP1s (D).

**Supplemental Figure 5:**
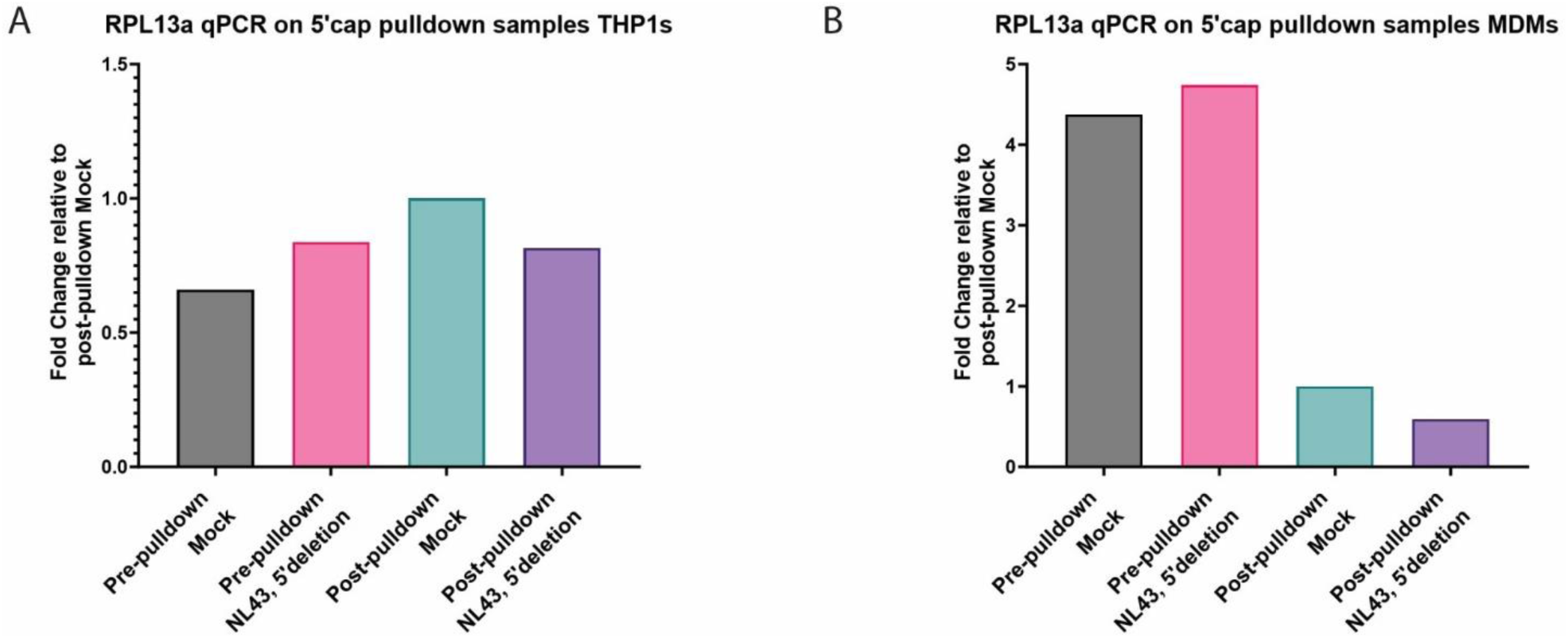
A-B) RTqPCR for RPL13a, a 5’capped mRNA transcript constitutively expressed in all cells, on RNA from MDMs (A) or PMA-differentiated THP1s (B) before selection for 5’cap mRNA or after selection for 5’cap mRNA.

**Supplemental Table 1:**
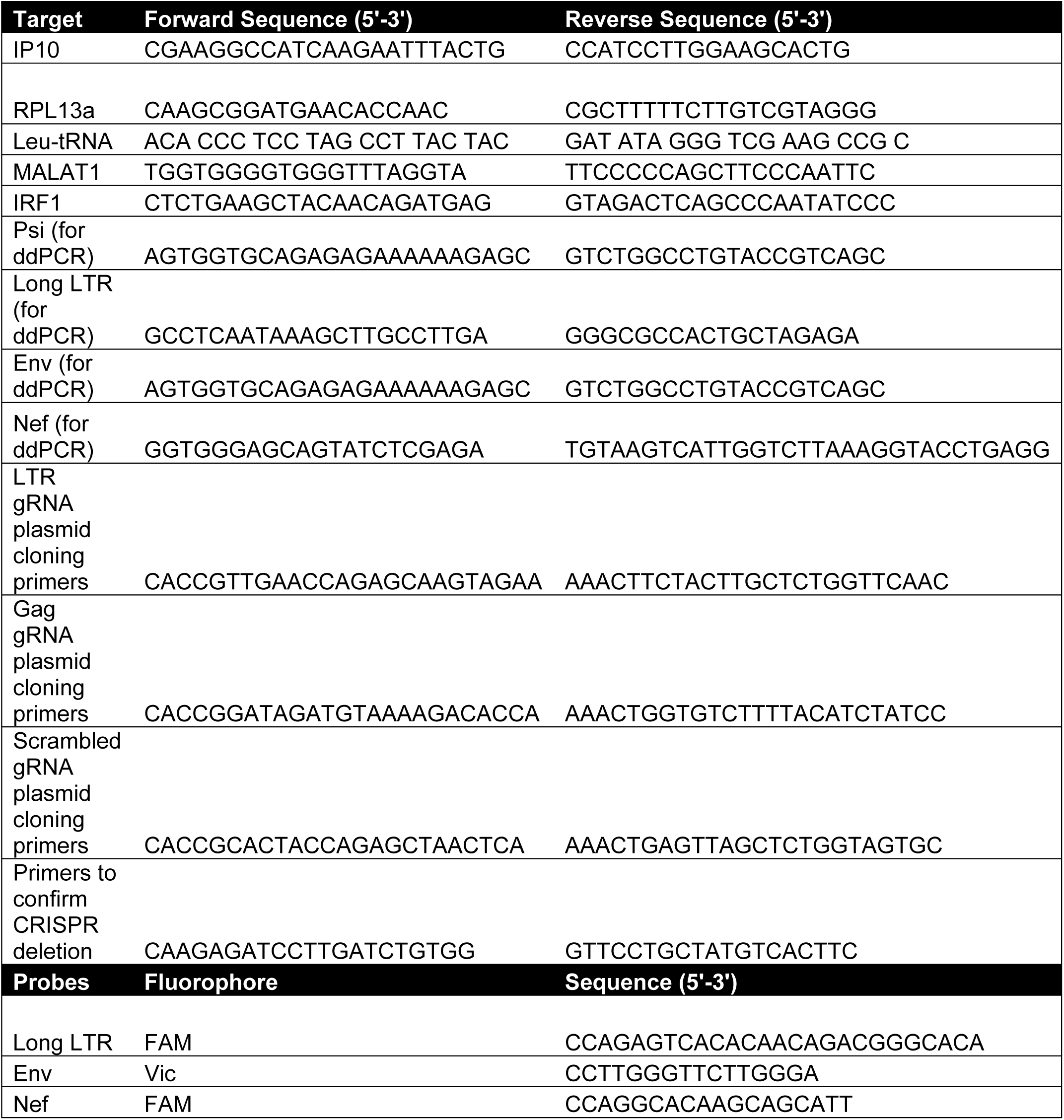
Primer and Probe Sequences.

**Supplemental Table 2:**
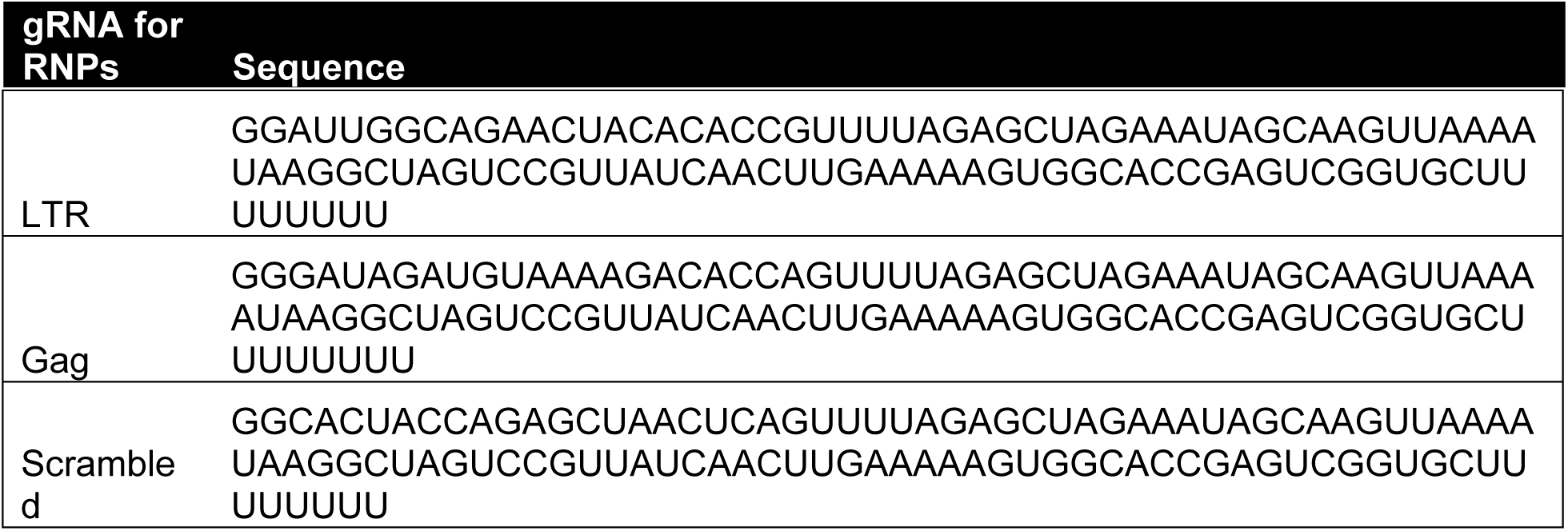
gRNA Sequences.

**Supplemental Table 3:**
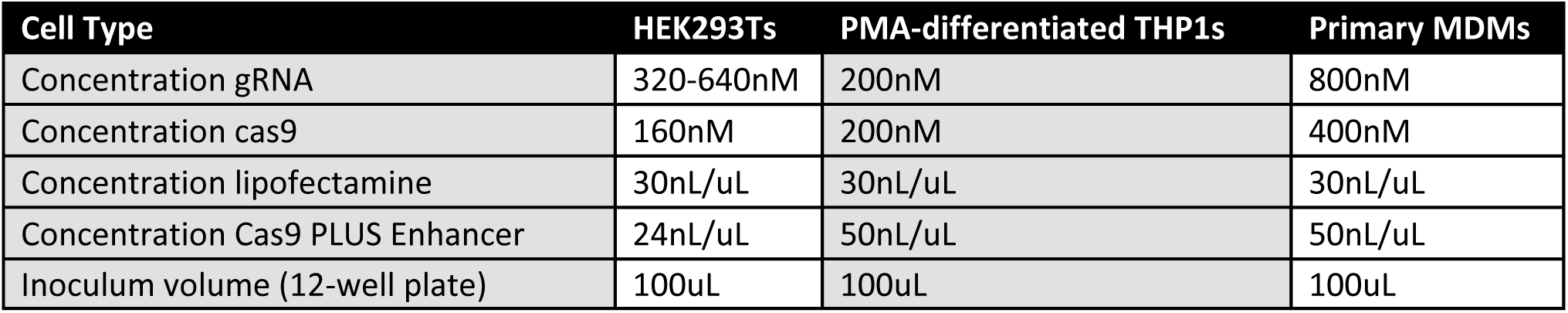
Concentration of RNP components in master mix.

| Supplemental Table 3: Concentration of RNP components in master mix |  |  |  |
| --- | --- | --- | --- |
| Cell Type | HEK293Ts | PMA-differentiated THP1s | Primary MDMs |
| Concentration gRNA | 320-640nM | 200nM | 800nM |
| Concentration cas9 | 160nM | 200nM | 400nM |
| Concentration lipofectamine | 30nL/uL | 30nL/uL | 30nL/uL |
| Concentration Cas9 PLUS Enhancer | 24nL/uL | 50nL/uL | 50nL/uL |
| Inoculum volume (12-well plate) | 100uL | 100uL | 100uL |

**Supplemental Table 4:**
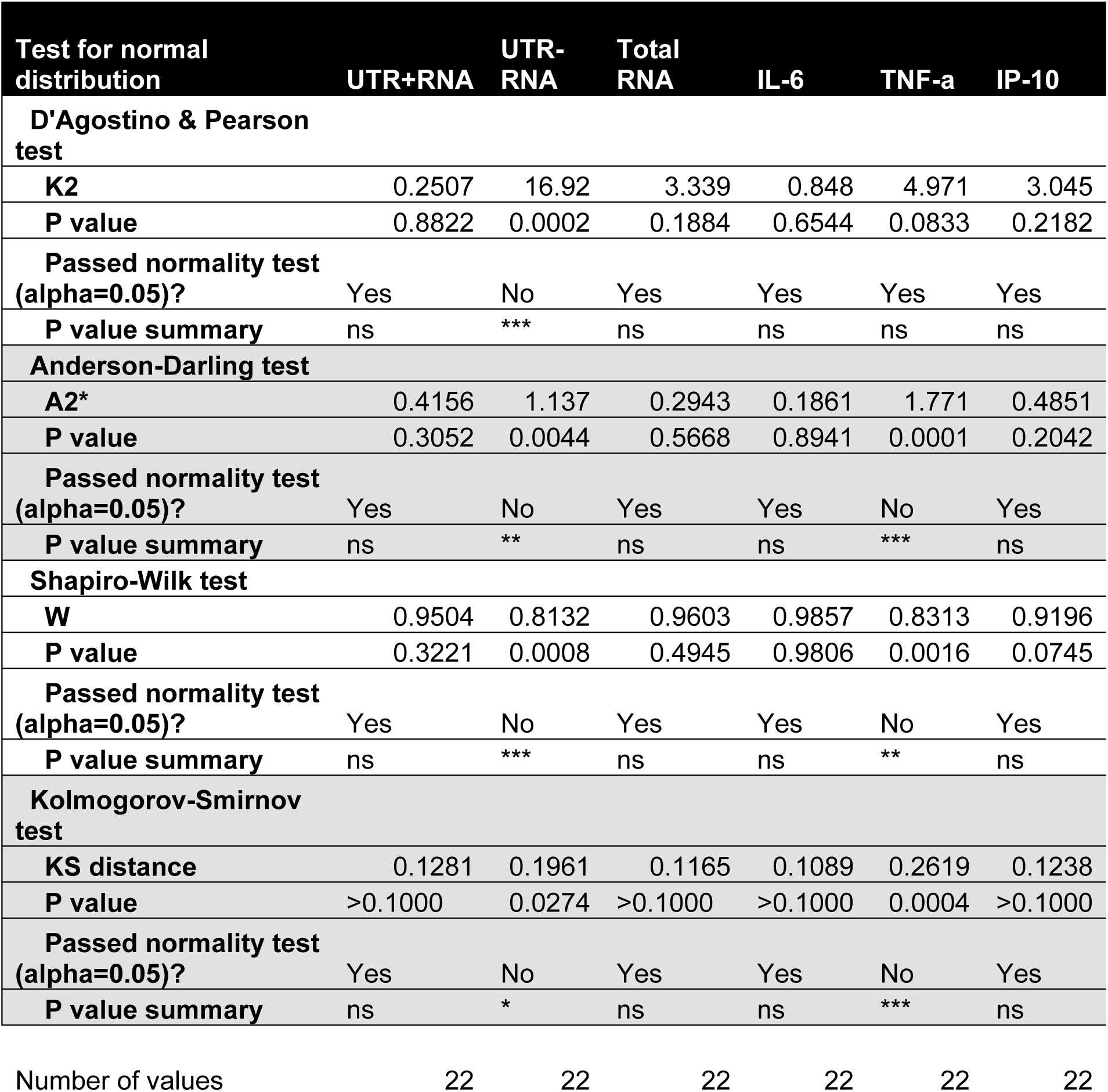
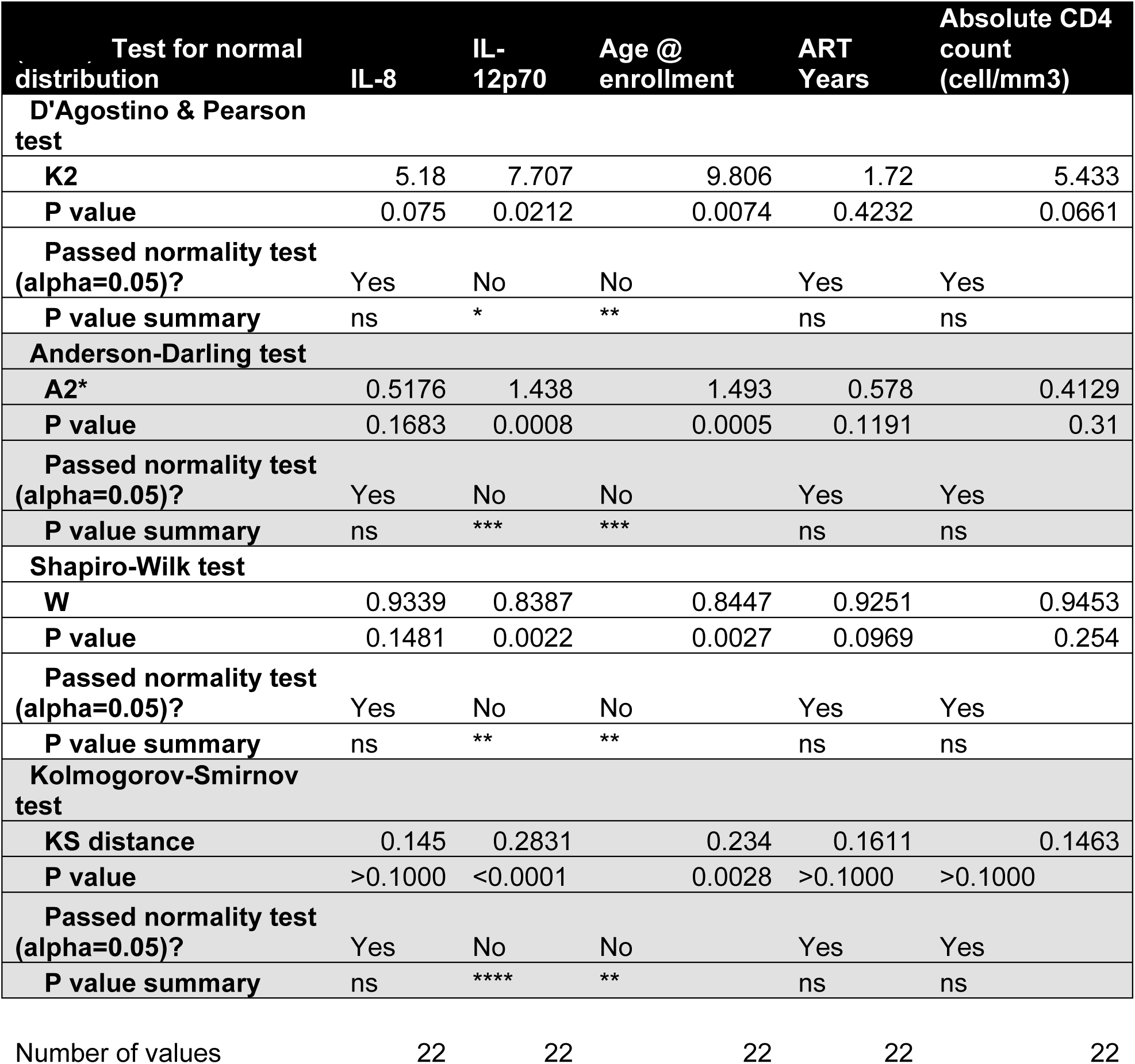

**Supplemental Table 5:** Multiple Linear Regression of cell-associated UTR-deficient RNAs vs IP10 in serum.

| Variable | $\beta$ | 95% CI (asymptotic) | P value | P value summary |
| --- | --- | --- | --- | --- |
| Intercept | 0.1235 | -0.5675 to 0.8145 | 0.7108 | ns |
| UTR-deficient RNA | 0.4581 | 0.03540 to 0.8807 | 0.0353 | * |
| Age @ enrollment | 0.007245 | -0.006340 to 0.02083 | 0.2762 | ns |
| ART Years | -0.004207 | -0.03632 to 0.02790 | 0.7855 | ns |
| Sex @ birth | -0.2098 | -0.5890 to 0.1695 | 0.2593 | ns |

**Supplemental Table 6:** Multiple Linear Regression of cell-associated UTR-deficient RNAs vs TNFα in serum.

| Variable | $\beta$ | 95% CI (asymptotic) | P value | P value summary |
| --- | --- | --- | --- | --- |
| Intercept | 1.167 | 0.6041 to 1.730 | 0.0004 | *** |
| UTR-deficient RNA | 0.4139 | 0.06950 to 0.7583 | 0.0213 | * |
| Age @ enrollment | 0.002356 | -0.008713 to 0.01343 | 0.659 | ns |
| ART Years | -0.004376 | -0.03054 to 0.02179 | 0.7285 | ns |
| Sex @ birth | 0.1226 | -0.1864 to 0.4316 | 0.4142 | ns |

**Supplemental Table 7:**
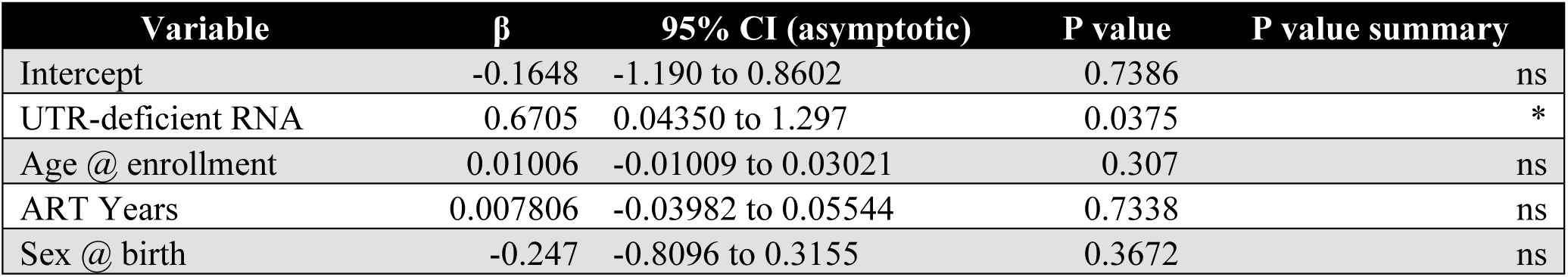
Multiple Linear Regression of cell-associated UTR-deficient RNAs vs IL-8 in serum.

| Variable | $\beta$ | 95% CI (asymptotic) | P value | P value summary |
| --- | --- | --- | --- | --- |
| Intercept | -0.1648 | -1.190 to 0.8602 | 0.7386 | ns |
| UTR-deficient RNA | 0.6705 | 0.04350 to 1.297 | 0.0375 | * |
| Age @ enrollment | 0.01006 | -0.01009 to 0.03021 | 0.307 | ns |
| ART Years | 0.007806 | -0.03982 to 0.05544 | 0.7338 | ns |
| Sex @ birth | -0.247 | -0.8096 to 0.3155 | 0.3672 | ns |

